# The neurocellular implementation of representational geometry in primate prefrontal cortex

**DOI:** 10.1101/2023.03.06.531377

**Authors:** Xiao-Xiong Lin, Andreas Nieder, Simon N. Jacob

## Abstract

Modern neuroscience has seen the rise of a population-doctrine that represents cognitive variables using geometrical structures in activity space. Representational geometry does not, however, account for how individual neurons implement these representations. Here, leveraging the principle of sparse coding, we present a framework to dissect representational geometry into biologically interpretable components that retain links to single neurons. Applied to extracellular recordings from the primate prefrontal cortex in a working memory task with interference, the identified components revealed disentangled and sequential memory representations including the recovery of memory content after distraction, signals hidden to conventional analyses. Each component was contributed by small subpopulations of neurons with distinct electrophysiological properties and response dynamics. Modelling showed that such sparse implementations are supported by recurrently connected circuits as in prefrontal cortex. The perspective of neuronal implementation links representational geometries to their cellular constituents, providing mechanistic insights into how neural systems encode and process information.

## Introduction

For decades, the dominant approach to understanding neural systems has been to characterize the role and contributions of individual neurons. In a recent paradigm shift, the concept of high-dimensional activity spaces that represent cognitive and other variables at the level of neuronal populations has taken the center stage and sidelined the single-neuron perspective (Barack & Krakauer, 2021; Saxena & Cunningham, 2019). These population representations capture multi-neuron activity in different behavioral task conditions in the form of geometrical structures (Bernardi et al., 2020; Okazawa et al., 2021). Representational geometry provides a complete description of the information encoded by and processed in a neuronal population. It does not, however, account for how individual neurons – the nuts and bolts of brain processing – give rise to the representations and the operations performed on them (Kriegeskorte & Wei, 2021) because there is no direct connection between informational representation and biological implementation at the cellular and circuit level.

In constructing representational geometries, the choice of coordinate system, that is the set of components that capture the population activity, is arbitrary. The question then arises what the most meaningful coordinate system is to represent the data. In principal component analysis (PCA), a widely used method for dimensionality reduction, the principal components (PCs) capture the neuronal activity’s variance, but they are not designed to yield biologically interpretable aspects of the representational geometry. Identifying coordinate systems that are rooted in biology is particularly relevant in association cortices where neurons often have mixed-selective responses that are not easily interpreted as the representation of any single stimulus or task variable alone (Bernardi et al., 2020; Rigotti et al., 2013). Neuronal signals in association cortices also show complex temporal dynamics and task-dependent modulations that reflect distinct sensory and memory processing stages (Cavanagh et al., 2018; Jacob et al., 2018; Jacob & Nieder, 2014). During working memory, for example, behaviorally relevant target items are maintained in online storage and must be protected against interfering distractors (Jacob et al., 2018; Jacob & Nieder, 2014). However, depending on which coordinate system is used to express the representational geometry, the same task-related neuronal activity could be interpreted in one of two ways: either as components representing the target in each task epoch individually, suggesting a memory mechanism built on sequential relay of target information among components (Parthasarathy et al., 2019), or, alternatively, as components that represent the target across task epochs, suggesting a memory mechanism of continuous representation of target information by the same components (Tang et al., 2020).

The biological implementation of representations points to how components are accessed and information is communicated. Unlike the units in neuronal network models, *in vivo* neurons are subject to anatomical and physiological constraints. There are approximately 10^10^ neurons in the human brain and 10^9^ in a hypothetical functional module such as the dorsolateral prefrontal cortex (PFC) (Courchesne et al., 2011; Herculano-Houzel et al., 2015). A pyramidal cortical neuron has on the order of 10^4^ dendritic spines (Eyal et al., 2018). Thus, given the disproportion between the low number of possible connections and the large number of potentially informative neurons, a neuron downstream of the PFC can only ‘read out’ from a small fraction of neurons in this region. That is, it cannot access arbitrary components of the representational geometry. Instead, it would be more efficient and biologically plausible to read out components that a few neurons predominantly contribute to, that is the components with a sparse neuronal implementation.

Here, we present a framework that exploits the structure in the representational geometry’s neuronal implementation. We show that this approach yields unbiased components of population activity that retain links to individual neurons. We performed data dimensionality reduction on extracellular multi-channel recordings from the non-human primate PFC by leveraging sparsity constraints in order to identify components that are contributed mainly by small subpopulations of strongly coding neurons (sparse component analysis, SCA; Georgiev et al., 2007; Olshausen & Field, 1996). We found that the activities on these components nontrivially matched the working memory task sequence performed by the animals, revealing separate sensory and memory components including a previously hidden component, namely the recovery of memory content after distraction. Notably, each component was made up of non-overlapping subpopulations of neurons with distinct electrophysiological properties and temporal dynamics. Finally, neuronal network modelling showed that recurrent connectivity as in the PFC favors such sparse implementations over non-structured Gaussian implementations. The framework and findings presented here bridge the gap between the single-neuron doctrine and the neuronal population doctrine (Barack & Krakauer, 2021; Saxena & Cunningham, 2019) and establish the perspective of neuronal implementation as an important complement to representational geometry.

## Results

### Different neuronal implementations may underlie the same representational geometry

Representational geometry abstracts the information coded by a population of neurons from their individual tuning profiles (Kriegeskorte & Wei, 2021). It specifies the pairwise distances between task-related collective neuronal responses, but no longer reflects the exact pattern of firing rates. This approach defines a stimulus-representing subspace. To illustrate, the representations for two stimuli A and B in PC space separate, rotate and collapse back to the origin (**Fig. 1a**).

**Fig. 1.**
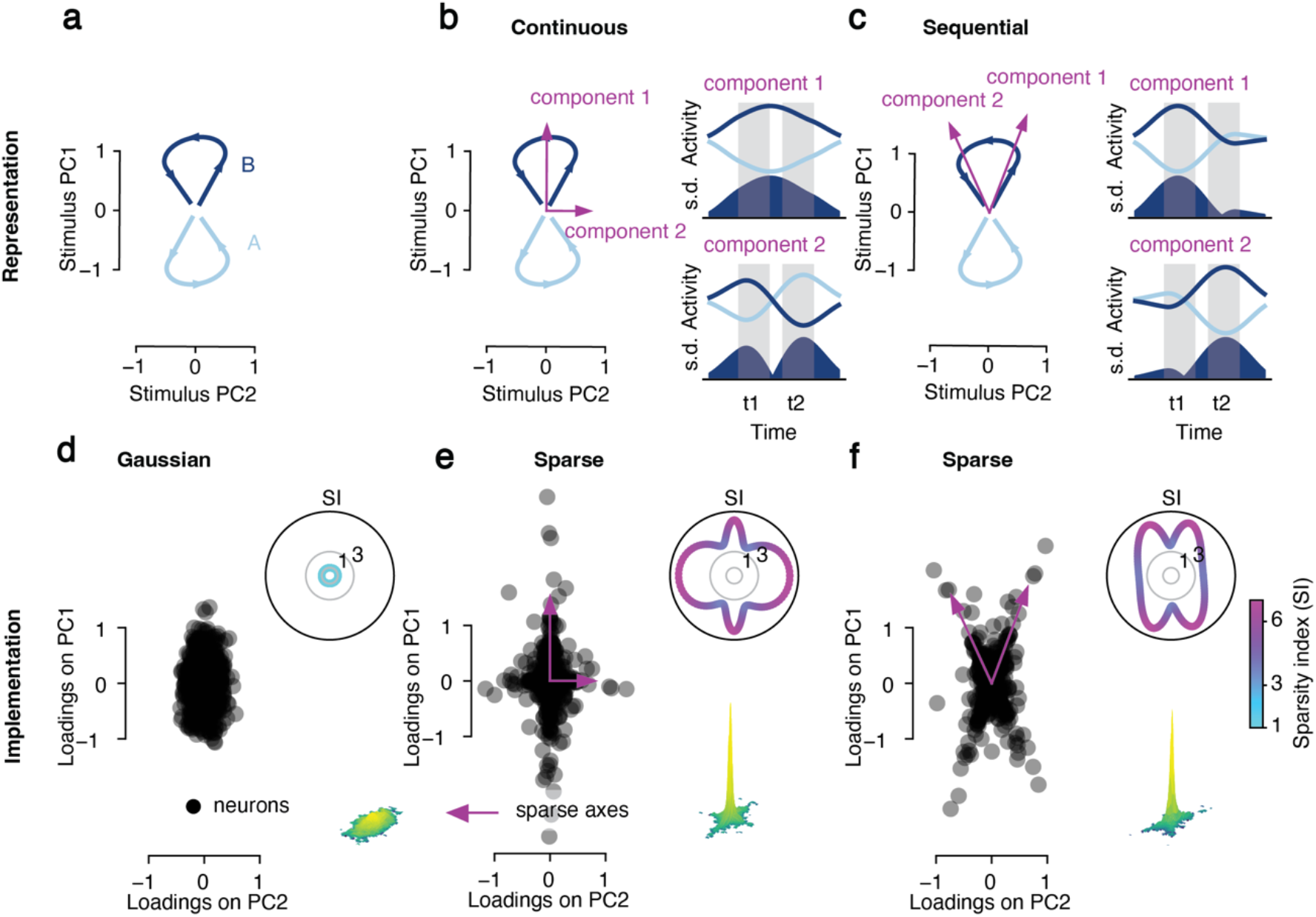
Different neuronal implementations of the same representational geometry. (**a**) Representational geometry for two trials with stimuli A and B on the plane specified by stimulus PC1 and PC2. Time runs along the individual trajectories. (**b**) Left: example pair of components that express the representational geometry (magenta arrows). Right: activities on the corresponding components and standard deviation (s.d.) across components as a measure of amount of information carried by them. Components are aligned with the PCs. (**c**) Same layout as in (b) for a non-aligned pair of components. (**d-f**) Neuronal implementation underlying the representational geometry in (a-c), specified by the distribution of neuronal loadings on the stimulus PCs. Insets: sparsity index (SI) of all axis orientations in the space spanned by PC1 and PC2. Axes with high SI (sparse axes, magenta arrows) in (e) and (f) correspond to the components 1 and 2 in (b) and (c), respectively.

The same stimulus-representing subspace can be defined with arbitrary sets of components. Components can be chosen to capture specific aspects of the representation, e.g., to continuously distinguish between stimuli (**Fig. 1b**), or to distinguish between stimuli at different time points (**Fig. 1c**). Note that in the former example, the components align with the PCs, while in the latter they do not. Various studies have followed this approach, selecting the components e.g. such that they express representations sequentially (Aoi et al., 2020) or such that they each correspond to a particular task variable of interest (Libby & Buschman, 2021; Mante et al., 2013).

Neuronal activity can be reconstructed by the weighted sum of components. Every neuron has a set of weights quantifying its relation to the different components, i.e. its loadings on the components. The loadings of neurons on the PCs visualize their positions in implementation space (**Fig. 1d-f**), where the loadings along any axis correspond to a component in representation space with the same orientation (**Fig. 1a-c**). The structure in the implementation space, i.e., the distribution of loadings across neurons, can be exploited to identify a unique, non-arbitrary set of components that emphasizes biological plausibility of stimulus coding over enforcing possibly unjustified priors.

Representational geometry is invariant to the rotation of neuronal coordinates (Kornblith et al., 2019). Different neuronal implementations may therefore underlie the same representational geometry. We first consider the scenario of a Gaussian (dense) distribution of loadings (**Fig. 1d**), where the standardized moments (e.g., skewness and kurtosis) are constant, meaning there are no differences in these distributional statistics across axis orientations. We define the sparsity index (SI; **Fig. 1d**, top inset) to denote the sparsity of the implementation along a given axis. SI is proportional to a distribution’s kurtosis. If SI is constant across axis orientations, neurons do not preferentially align to any axes.

Next, we consider a sparse distribution (**Fig. 1e**). Most neurons lie around the origin of the coordinate system. However, because SI is not constant (**Fig. 1e**, top inset), we can find the sparse components that strongly coding neurons align to. In the present case, these sparse axes correspond to the components in representational space that code the difference between stimulus A and B continuously (with one of the components reversing between epochs; compare **Fig. 1e** with **Fig. 1b**). Importantly, sparse distributions can exist for arbitrary axis orientations. For example, strongly coding neurons could align to the components that sequentially represent the stimulus information at time point 1 and time point 2 (compare **Fig. 1f** with **Fig. 1c**).

Although both scenarios are characterized by sparse neuronal implementations, we note that they have fundamentally different implications for readout, lending particular importance to the positioning of sparse axes orientations. Continuous readout (**Fig. 1b** and **e**, component 1) is stable, but not optimized for either time point 1 or time point 2, whereas sequential readouts (**Fig. 1c** and **1f**) are more precise at the respective time points, but not stable across time points.

In summary, the perspective of neuronal implementation offers a way to connect representational geometries to their cellular constituents, revealing mechanistic insights into how a neural system encodes, processes and relays information.

### The neuronal implementation of working memory

With this framework, we now examine neuronal implementation of working memory, a core cognitive function for online maintenance and manipulation of information in the absence of sensory inputs. Extracellular multi-channel recordings were performed in the lateral PFC of two monkeys trained on a delayed-match-to-numerosity task, requiring them to memorize the number of dots (i.e., numerosity) in a visually presented sample and resist an interfering distracting numerosity (Jacob and Nieder, 2014) (**Fig. 2a**). A total of 467 single units recorded across 78 sessions were included in the analysis. Spike rates were binned, averaged across conditions of the same type and demixed into their constituent parts (**Fig. 2b**) (Kobak et al., 2016). Because the task design was balanced (i.e., all sample-distractor combinations were included), the different task variables were statistically independent of each other. Demixing therefore allowed to isolate and analyze signal components that would otherwise be overshadowed by signals that dominate the raw firing rates. Across neurons, the neuronal activities coding for trial time, sample numerosity, distractor numerosity and the sample-distractor interaction accounted for 72.7 %, 8.7 %, 5.8 % and 12.9 % of the total variance, respectively (**Fig. 2b**).

**Fig. 2.**
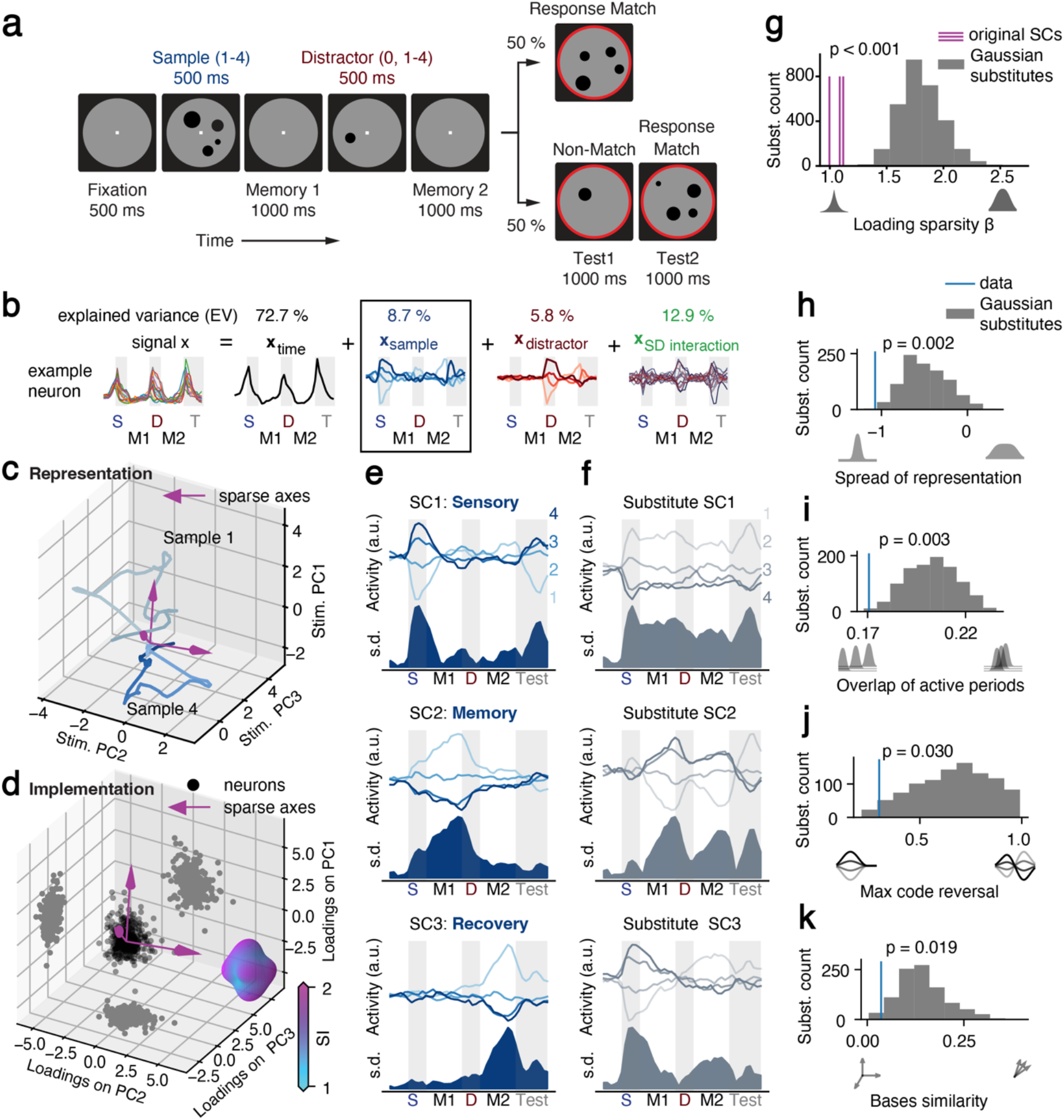
The neuronal implementation of working memory. (**a**) Delayed-match-to-numerosity task with distractors. (**b**) Demixing procedure separating the activity of each neuron into the parts coding time, sample numerosity, distractor numerosity and sample-distractor interaction. The sample coding part is used for the following analyses. Top: percentage of explained variance for each part. (**c**) Representational geometry for sample numerosities 1 and 4 in PC space, averaged across trials of the same condition. (**d**) Loadings of all recorded neurons on the top three PCs (black dots) including distributions projected onto the planes formed by PC pairs (gray dots). Sparse axes (magenta arrows; determined by SCA) have high SI. Inset: surface plot of SI for all axes in the space. (**e**) Activity of the three identified sparse components (SCs), averaged across trials for each sample numerosity condition (top; numbers indicate sample numerosity) and relative information across conditions measured as standard deviation (s.d.). (**f**) SCs of an example substitute dataset with non-structured Gaussian implementation. (**g**) Sparsity β of the neuronal loadings on the SCs (fit to generalized normal distribution) for the original data and the substitute datasets (permutation test with n = 3×1000 permutations). (**h-k**) Activity measures for the SCs of the original data and the substitute datasets (permutation test with n = 1000 permutations).

We first focused on the representation of the sample numerosity throughout the trial, the crucial function for completing the task (**Fig. 2c**). In PC space, the representations of different numerosities (1 and 4 visualized here) started to separate, marking an increase of the information during sample presentation. Then the representations rotated and returned to the origin. Similar representational changes have been reported previously (Elsayed & Cunningham, 2017; Murray et al., 2017; Parthasarathy et al., 2019).

The distribution of loadings of individual neurons onto the first three PCs was highly non-Gaussian (p < 0.001; Henze-Zirkler multivariate normality test; **Fig. 2d**). Accordingly, the sparsity index (SI) was not uniform across all axis orientations (**Fig. 2d**). Using sparse component analysis (SCA) that identifies components with sparse distributions of neuronal loadings (sparse components, SCs), we found three SCs that optimally decomposed the sample numerosities’ representational geometry. The SCs displayed temporally well-defined active periods that matched the task structure and tiled the duration of a trial (**Fig. 2e**). Intuitively, they correspond to components for sensory encoding, memory maintenance and memory recovery following distraction, in accord with the scenario of sequential representations (cp. to **Fig. 1c** and **f**).

To control for the possibility that noise in non-sparse implementations is mistaken for structure by SCA, we created substitute datasets with random Gaussian implementations (i.e., Gaussian distributions of neuronal loadings) while keeping the representational geometry intact and then systematically compared the original SCs with the substitute SCs (example substitute SCs in **Fig. 2f**). First, the sparsity parameter (fit to the distribution of loadings on the SCs) was smaller for all three original SCs than for the substitutes (p < 0.001 for all three SCs; permutation test with n = 3×1000 permutations; **Fig. 2g**), confirming the presence of structure in the implementation. Second, the activities on the SCs showed temporally restricted sample representations with shorter spread (p < 0.002; permutation test with n = 1000 permutations; same as for Fig. 2i-k; **Fig. 2h**), less temporal overlap with other SCs (p < 0.003; **Fig. 2i**), and less reversal of sample numerosity tuning (p < 0.030; **Fig. 2j**) than the substitutes, suggesting that the observed SC activity was more sequential than to be expected with a random implementation. Third and finally, the SCs were closer to orthogonal than the substitutes (p < 0.019; **Fig. 2k**), demonstrating that the observed implementation is more efficient than a random implementation.

In summary, the neuronal implementation of the sample numerosities’ representational geometry was structured and sparse. The activities on the sparse components demonstrated sequential rather than continuous coding of working memory content, indicating that the change of behavioral demands in the course of the trial triggers a switching of informative subpopulations.

### The effect of distraction on sample numerosity representations

The lack of a component that continuously represented the behaviorally relevant sample numerosity throughout the trial was unexpected. We therefore investigated the influence of distraction on sample number coding.

First, we applied SCA to the demixed distractor coding part of the data (**Fig. 3a**, top). Two SCs were obtained that were sequentially active during presentation and maintenance of the distractor numerosity, respectively (**Fig. 3a**, bottom). These components resembled the sensory and memory sample coding SCs (cp. to **Fig. 2e**), suggesting that target and distracting information initially occupied similar resources despite their distinct behavioral relevance. Supporting this hypothesis, we found strongly overlapping neuronal loadings between sample SCs and distractor SCs (cosine similarity; 0.69 and 0.57 for the sensory and memory components, respectively; **Fig. 3b**) with displacement of sample information by distractor information as the trial evolved (**Fig. S1a**, top and middle). However, in contrast to the sample sensory and memory components, the sample recovery SC was unique and did not share loadings with any other SC (Fig. 3b). Furthermore, the sample recovery SC was not influenced by distractor information and carried sample information until test numerosity presentation (**Fig. S1a**, bottom). To correctly complete a trial, more activity in the sample sensory and recovery SCs was required when the trial contained a distractor than when a trial without a distractor was presented (**Fig. S1b**). Conversely, distractors led to reduced sample activity in the memory component.

**Fig. 3.**
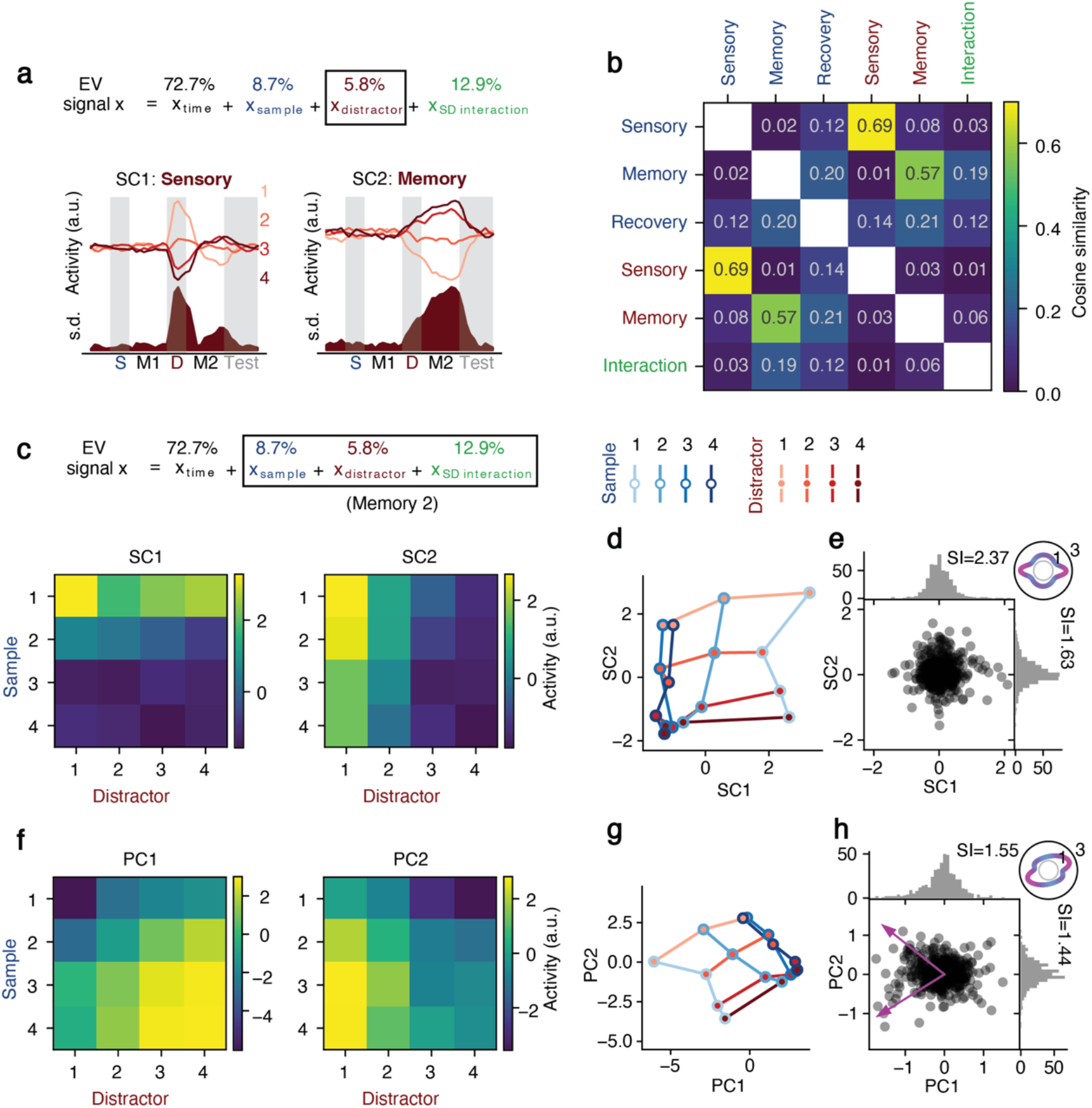
The effect of distraction on sample representations. (**a**) Top: the demixed distractor representing part used in the analysis. Bottom: distractor numerosity sparse components (SCs). Numbers indicate distractor numerosity. (**b**) Cosine similarity between loadings of sample numerosity SCs (blue), distractor numerosity SCs (red) and the sample-distractor interaction SC (green). (**c**) Activity of the two SCs identified using firing rates averaged across the second memory delay for all sample-distractor combinations without demixing the stimulus presentations. (**d**) Representational geometry in SC space. Blue and red colors indicate sample and distractor numerosity, respectively. (**e**) Neuronal loadings on the 2 SCs. Dots: joint distribution in SC space. Histograms: marginal distribution of neuronal loadings on SC1 and SC2. Inset: SI for all axes. (**f-h**) Same layout as in (c-e) but for PCs. Magenta arrows in (H) indicate sparse axes.

Second, we applied SCA to the sample-distractor interaction part of the data. One SC was identified. Its activity was most pronounced when the sample and distractor numerosity were the same (**Fig. S2**). The neuronal loadings on this SC did not overlap with the loadings on sample or distractor SCs (**Fig. 3b**), suggesting that the boost in numerosity information was generated by a dedicated subpopulation responding to a repeated presentation of the same number, instead of changing the activity of the sample representing neurons.

Together, these results indicate a (partially) shared capacity for sample and distractor representations during the sensory input and subsequent memory delay stages. The invasion of distractor information forced the recruitment of an extra component, the recovery component, to maintain sample information in working memory.

So far, all analyses were performed on separated (demixed) representations. We next investigated whether sample and distractor information could be equally disentangled using SCA alone without demixing the numerosity coding signal (**Fig. 3c**). SCA performed on firing rates averaged across the second memory delay recovered two sparse components that each selectively captured sample and distractor information (**Fig. 3d**). The corresponding representational geometry was grid-like with clearly factorized sample and distractor information that each aligned well to one SC (**Fig. 3e**). Notably, this alignment was non-trivial and not enforced by our analytical method, arguing that the PFC spontaneously disentangles target and distractor representations in working memory. The underlying implementation showed clear sparse structure in the neuronal loadings onto these components (**Fig. 3f**).

For comparison, PCA, which is insensitive to the neuronal implementation, was unable to recover factorized components (**Fig. 3g**). The grid-like geometry was still largely preserved, but it did not align with the PCs (**Fig. 3h**). In contrast to SCA, PCA did not identify the components with the sparsest loadings (**Fig. 3i**).

### Subpopulations of neurons dominate working memory representations

Next, we investigated whether the implementation was sparse enough to be able to reliably reconstruct the population-level sample representation using only a small fraction of neurons. We performed cross-temporal linear discriminant analysis (LDA) to decode sample numerosity at a given time point in the trial using training data from a different time point (**Fig. 4**). Decoding accuracy therefore quantifies the degree to which the representation is transferable. With four numerosities, chance level accuracy is 25 %. Using the entire population of 467 recorded neurons, we found a highly dynamic code with good within-epoch transfer, but very little generalization across epochs, in particular from the first to the second memory delay (**Fig. 4a**). In line with our previous results, this finding suggests that working memory representations are non-uniform and that distinct, complementary processes are required to protect behaviorally relevant information from interference.

**Fig. 4.**
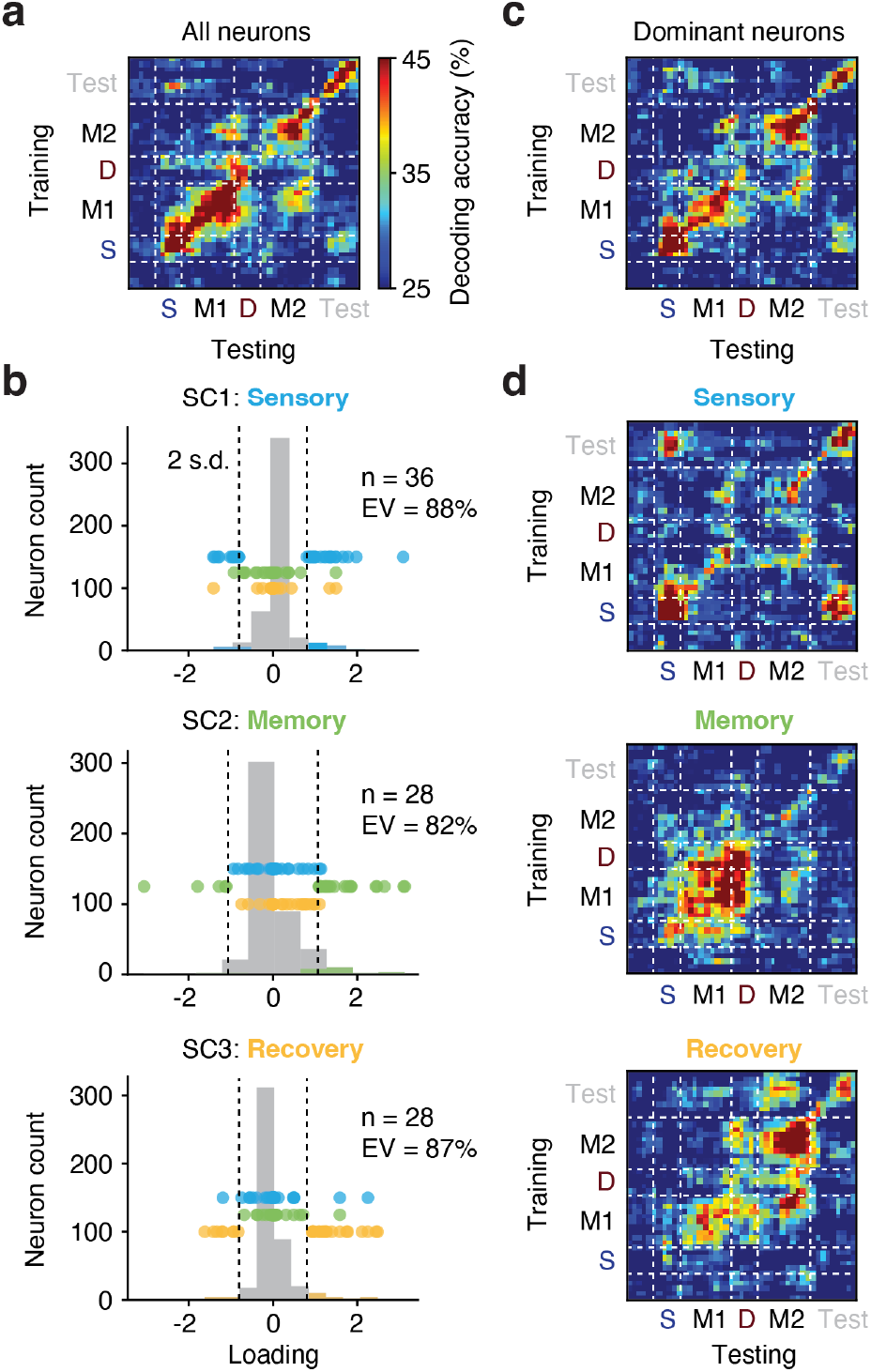
Subpopulations of neurons dominating working memory coding. (**a**) Accuracy of cross-temporal linear discriminant analysis (LDA) decoding of sample numerosity using all recorded neurons (y axis: training, x axis: testing). (**b**) Neuronal loadings on the three identified sample numerosity SCs. Colored dots indicate the ‘dominant’ neurons selected in each SC (cut-off: two s.d.). The percentage of variance explained within each SC is given for each subpopulation. (**c**) Accuracy of cross-temporal LDA decoding of sample numerosity using only the dominant neurons. Compare to (a). (**d**) Sample numerosity decoding accuracy using the dominant subpopulations of each SC. Same color scale in (a), (c) and (d).

We selected the neurons that contributed most to the previously identified SCs (loading on the SC larger than two standard deviations; **Fig. 4b**). 36, 28 and 28 single neurons passed the criterion for the sensory, memory and recovery SC, respectively. Although each subpopulation comprised only 6 to 8 % of the entire recorded population, these ‘dominant neurons’ explained 88 %, 82 % and 87 % of their respective component’s variance (sum of squares of dominant neurons’ loadings over sum of squares of all neurons’ loadings). Overlapping membership in two subpopulations was very rare (no more than three neurons in any SC pair; **Fig. 4b**).

Cross-temporal LDA using only the dominant neurons showed a very similar sample numerosity decoding pattern as with the entire population (**Fig. 4c**, cp. with **Fig. 4a**), confirming that the decoder previously relied mainly on this small subset of neurons. The sensory subpopulation contributed to decoding in particular during the sample and test numerosity presentation, but showed very little activity in the memory epochs (**Fig. 4d**, top). The memory subpopulation dominated in the first delay, but surprisingly was not involved in sample coding during the second delay (**Fig. 4d**, middle). Instead, after distraction, the recovery subpopulation was exclusively responsible for carrying sample information (**Fig. 4d**, bottom). This suggests that these neurons crucially contribute to shielding working memory information from interference (see also **Fig. S1**).

### Subpopulation-specific electrophysiological properties

Above, we identified dominant neurons based on their stimulus selectivity. We now investigated whether their different roles in representing sample information were possibly mirrored by distinct electrophysiological properties.

First, we calculated the across-trial similarity (Pearson correlation) between each neuron’s activity at different time points in the fixation period in order to derive the intrinsic time scale, a measure considered to index a neuron’s ability to maintain memory traces (Murray et al., 2014). Representative neurons from all three subpopulations are shown (**Fig. 5a**). The example recovery neuron had a significantly larger spread from the diagonal than the sensory and memory neuron, i.e., its activity in distant time points was more strongly correlated, thus signifying a longer time constant (**Fig. 5a**, bottom panel). For each subpopulation, an exponential decay was fitted to the mean correlation coefficient across neurons (**Fig. 5b**). The recovery subpopulation had the largest time constant τ (165 ms, 127 ms, and 338 ms for sensory, memory and recovery neurons, respectively). The distribution of τ values in the recovery population also stood out from the distributions observed in subsampled subpopulations of PFC neurons, whereas the sensory and memory neurons’ distributions were not significantly different (p = 0.874, p = 0.455, p = 0.002 for sensory, memory and recovery subpopulations, respectively; KL-divergence with bootstraps; **Fig. 5c**).

**Fig. 5.**
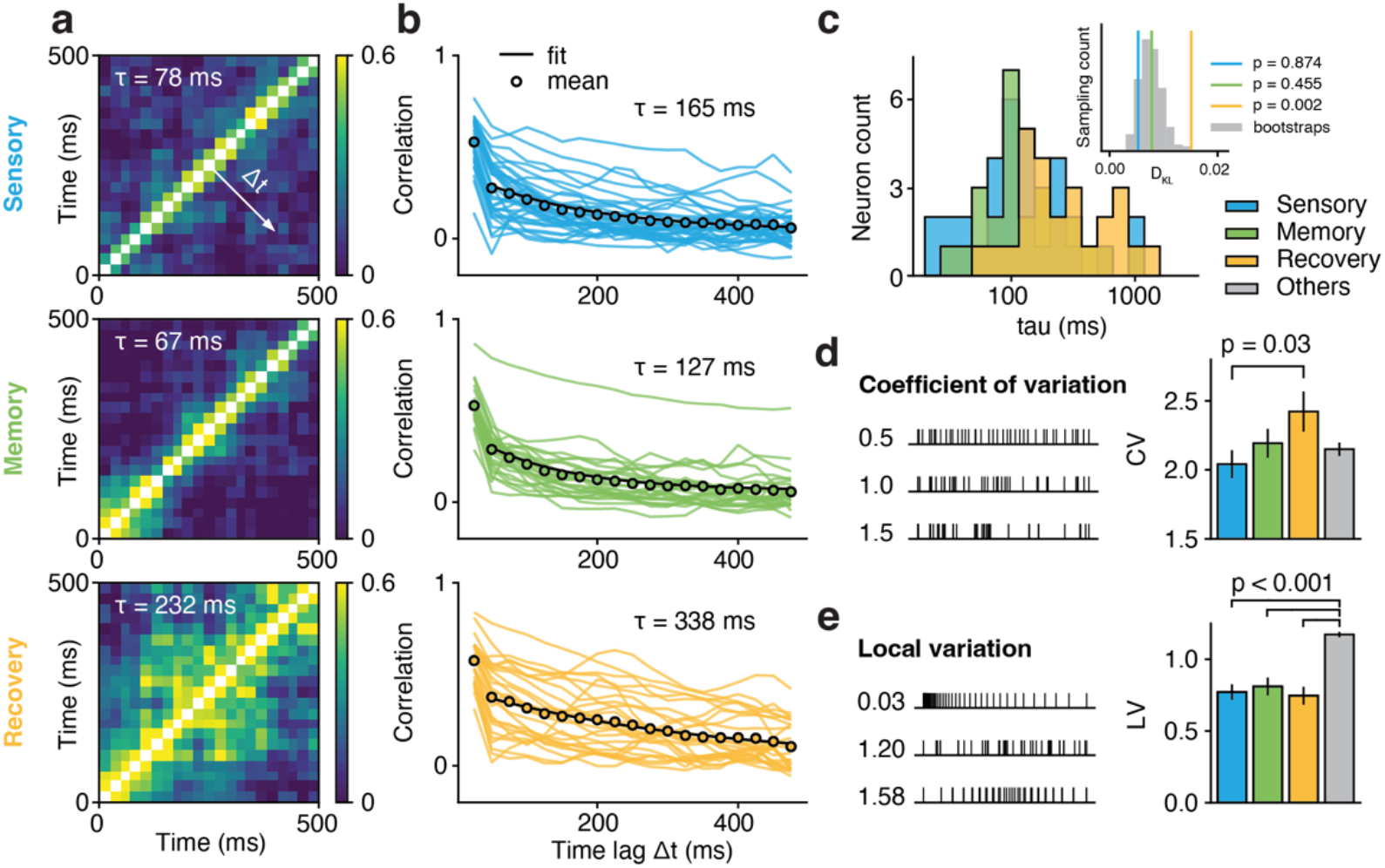
Subpopulation-specific electrophysiological properties. (**a**) Between-timepoint Pearson correlations of the trial-to-trial fluctuation of firing rates in the fixation epoch for the three dominant subpopulations. (**b**) Auto-correlograms obtained by averaging across diagonal offsets in (a). Auto-correlograms of individual neurons are given (single lines) together with the subpopulation average and the fitted exponential decay (black dots and line, respectively). (**c**) Distribution of fitted decay constants of individual neurons in each dominant subpopulation. Inset: Kullback-Leibler divergence (D_KL_) between the distribution of each subpopulation and the whole population (null distribution for significance testing created with n = 1000 bootstraps from the whole population). (**d**) Coefficient of variation (CV) of inter-spike intervals (ISI) of the dominant subpopulations and the non-dominant other neurons (two-tailed *t*-Test). Left: example spike trains for different CVs. (**e**) Same layout as in (d) for the local variation (LV) of ISI.

Next, we investigated spike train statistics using the inter-spike intervals (ISI) measured during the neurons’ entire recording lifetime. The coefficient of variation (CV) measures the irregularity of a spike train (**Fig. 5d**). CVs of all recorded neurons were larger than 1 (i.e., more irregular than a Poisson process) with a gradual increase of spiking irregularity across the sensory, memory and recovery subpopulations. CVs in the recovery neuron population were significantly larger than in the sensory subpopulation (p = 0.030, two-tailed *t*-Test; **Fig. 5d**). The local variation (LV) measures local ISI differences and complements CV, which is a global measure. LVs in all dominant neurons were smaller than 1 (i.e., less local variation than a Poisson process) and significantly lower than in the non-coding PFC population (p < 0.001, two-tailed *t*-Tests; **Fig. 5e**).

Notably, these distinct electrophysiological properties were not involved in the original selection of subpopulations and therefore lend support to the notion that the implementation structure carries biological meaning.

### Subpopulation-specific temporal dynamics and representation of context

There was no perceptual cue in the working memory task specifying the difference between sample and distractor. This forced the animals to internally keep track of a trial’s temporal evolution. To investigate whether temporal dynamics and context played a role in supporting the subpopulation-specific stimulus representations, we next analyzed the temporal part of the demixed signal and visualized condition-averaged activity trajectories in each of the dominant subpopulations (**Fig. 6a**).

**Fig. 6.**
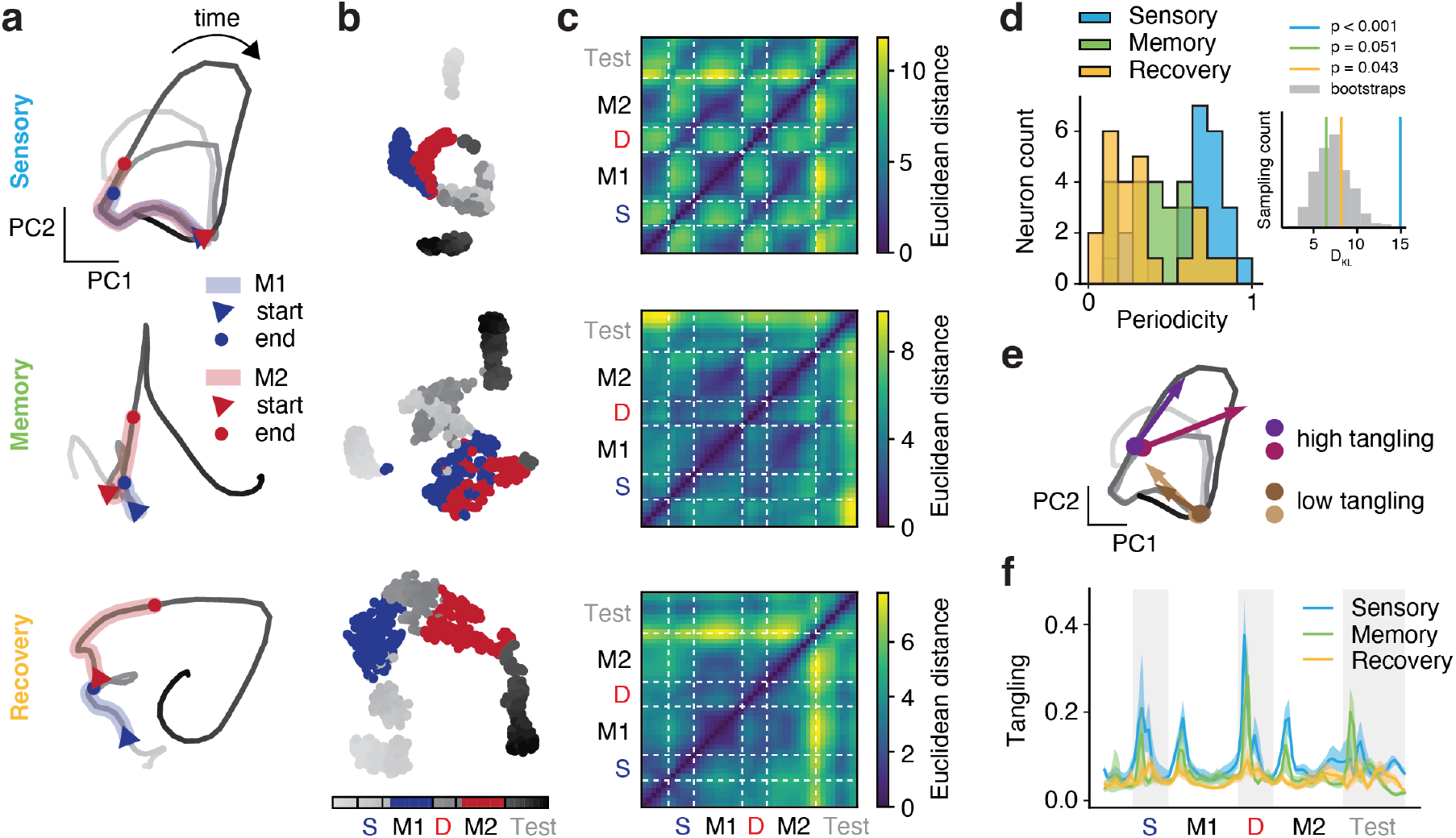
Subpopulation-specific temporal dynamics. (**a**) Temporal part of the demixed neuronal activity, averaged across conditions, of each dominant subpopulation projected onto their respective top two PCs. Time runs along the individual trajectories (bin width 50 ms). First and second memory delay are marked in blue and red, respectively. (**b**) Full signal averaged within each condition and embedded in 2D t-SNE space. Bins as in (a). (**c**) Euclidean distances between timepoints on the trajectory in (a) of each subpopulation. (**d**) Distribution of periodicity (relative power of 1/1.5 Hz and harmonics) of individual neurons in each subpopulation. Inset: Kullback-Leibler divergence (D_KL_) between the distribution of each subpopulation and the whole population (null distribution for significance testing created with n = 1000 bootstraps from the whole population). (**e**) Example timepoints on the trajectory of the sensory subpopulation with high and low tangling. **(f**) Time resolved tangling of the trajectory of each subpopulation.

In the sensory subpopulation, the trajectory followed a periodic, quasi-circular course (**Fig. 6a**, top panel). The first and second memory epochs overlapped almost entirely. This indicates that the sensory neurons did not distinguish between the time periods after sample and after distractor presentation. The trajectory of the memory subpopulation was less periodic, but intertwined in the first and second memory epochs (**Fig. 6a**, middle panel). In contrast, the trajectory of the recovery subpopulation was less intertwined, with most time points distinguishable from each other, especially the first and second memory epochs, signifying a better representation of the contextual difference following sample and distractor presentation (**Fig. 6a**, bottom panel).

Overlap of the memory epochs in the sensory and memory subpopulations could be due to the limitations of a linear projection and the emphasis of PCA on global structure. We therefore performed non-linear embedding using t-SNE (**Fig. 6b**). This analysis revealed comparable structures as the linear projection, with the first and second memory epochs separated only in the recovery neuron subpopulation.

To further investigate the temporal evolution of neuronal activity, we measured the Euclidean distances between individual time points in each subpopulation (full space; **Fig. 6c**). All distance matrices displayed a strong diagonal, reflecting the fact that close-by time points were represented similarly. Notably, there were also strong offset diagonals in the sensory subpopulation, meaning that activity in these neurons repeated with a cycle of about 1.5 s. Furthermore, activity in the sensory and memory epochs differed most in this subpopulation. These patterns were present, albeit weaker, in the memory subpopulation, but absent in the recovery neurons. We quantified periodicity for each neuron by computing the relative power of 1/1.5 s (0.67 Hz) activity and its harmonics normalized to the power of the full frequency spectrum (**Fig. 6d**). Compared to randomly sampled subpopulations of PFC neurons, the sensory subpopulation and the recovery subpopulation showed significantly different (higher and lower, respectively) periodicity (p < 0.001, p = 0.051, p = 0.043 for sensory, memory and recovery subpopulations, respectively; KL-divergence with bootstraps; **Fig. 6d** inset).

Neuronal activity is not static and temporally independent. Instead, firing rates at every time point depend on previous time points. To characterize the dynamical properties of the recorded PFC population in more detail, we used the measure of tangling (Russo et al., 2018). Tangling measures the extent to which the velocity (direction and speed) of a given state on a trajectory diverges from the velocity of its neighboring states (**Fig. 6e**), reflecting the level of unpredictability and instability (chaos) in the system. High tangling means a small disturbance in the current state would lead to large changes in the next state (difference of derivatives of neighboring points). The instability or inability to determine the next state from the current state (i.e., high tangling) indicates that other neuronal populations or external stimuli may drive the trajectory. Consequently, tangling was increased following the onset and offset of sensory input in all three subpopulations. Tangling was highest, however, in the sensory subpopulation and lowest in the recovery subpopulation (sensory vs. memory, p < 0.001; memory vs. recovery, p = 0.013; two-tailed *t*-Test across all trial time points; **Fig. 6f**).

In summary, these results suggest that the subpopulation of recovery neurons keeps a record of time and temporal context, which could contribute to these neurons’ ability to separate sample and distracting information. In contrast, the sensory subpopulation - and the memory subpopulation to a lesser degree - is characterized by its strong input-driven temporal dynamics, which is consistent with these neurons’ passive representation of numerosity regardless of it being behaviorally relevant (sample) or irrelevant (distractor).

### Recurrent connectivity favors sparse implementations

The implementation underlying the temporal evolution of neuronal representations is not arbitrary, but must be derived from the dynamical system of constituent neurons and their anatomical connectivity pattern. The PFC is a highly recurrent, rather than purely feed-forward, brain region (Harris et al., 2019). If biological structure and resource efficiency indeed favor sparse implementations, these should be better captured by recurrently connected networks than non-structured Gaussian implementations.

To address this hypothesis, we constructed a recurrent neural network model (RNN) to reproduce the target (to-be-fitted) firing rate sequences of each sample-distractor combination (**Fig. 7a**). The model consists of 467 neurons (to match the recorded population) receiving inputs of stimulus information according to the task structure. The model learns the recurrent connectivity *W* among the neurons. *W* summarizes the influence of the current time point’s firing rates ***r*** on the firing rates of the next time point. An indicator vector ***n*** (one non-zero entry) represents the sample and distractor numerosity, activating the numerosity-specific input in *I* to the entire neuronal population. To reflect the absence of an explicit visual cue that differentiates between sample and distractor in the task design, sample and distractor numerosity share the same input channel (*I*, ***n***). The contextual difference is left for the model to resolve. The intercept term ***b*** captures the baseline activity of each neuron.

**Fig. 7.**
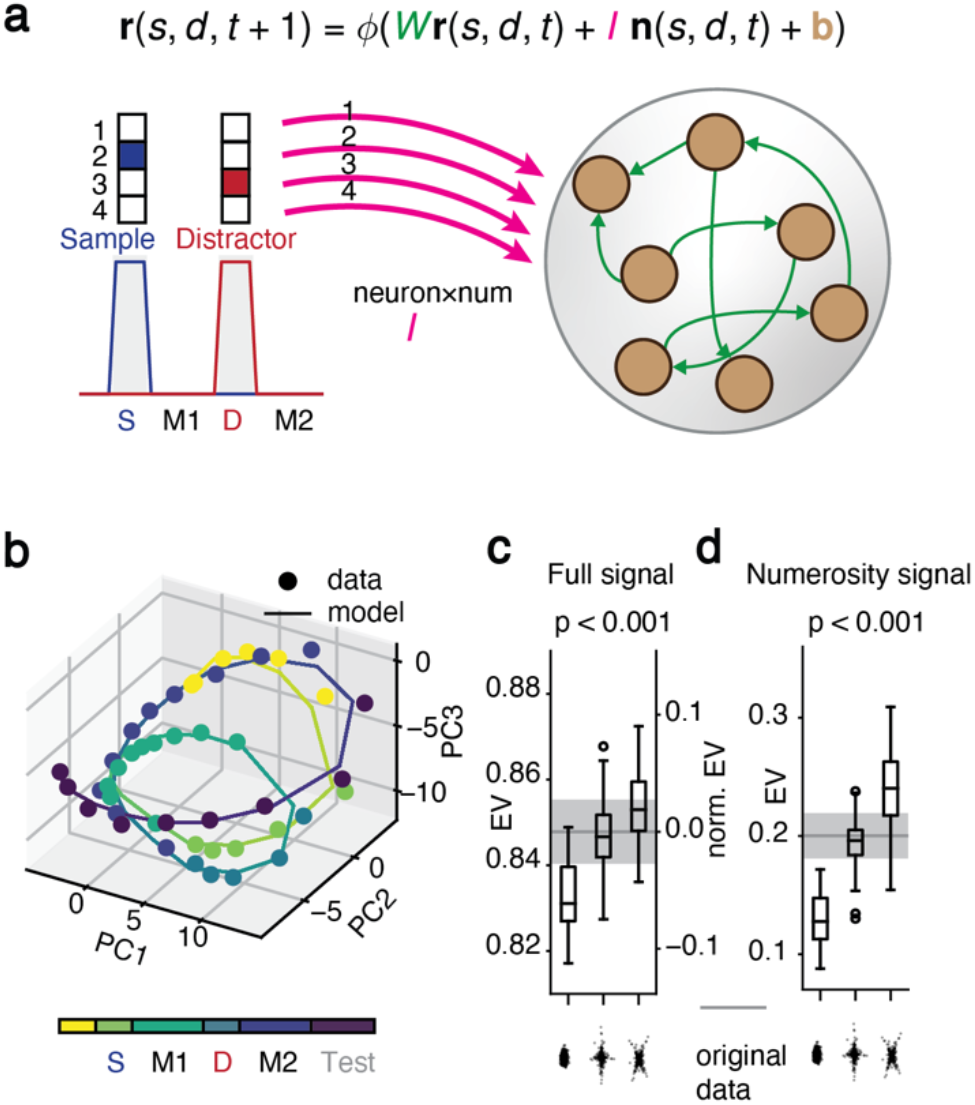
Recurrent neural network modeling. (**a**) RNN model governing equation and structure. Magenta and green arrows indicate numerosity-specific inputs and connectivity weights to be trained, respectively. (**b**) Model fit (solid trajectory) to original data (dots) averaged across all conditions. (**c**) Percentage of variance of the full signal explained by the model for non-structured Gaussian implementations of numerosity representations (left bar), sparse implementations with random orientations of sparse axes (middle bar) and sparse implementations with the same orientation of sparse axes as in the original data (right bar). Left and right axis show explained variance relative to the full signal and to the manipulated signal, respectively (one-way ANOVA across substitutes). (**d**) Same layout as in (c) for the percentage of variance of the numerosity signal explained by the model.

We first trained the model on the original dataset and visualized the trajectory of the output averaged across all conditions (**Fig. 7b**). The model reproduced the original dataset well, capturing 85.7 % of total variance. Next, we created substitute datasets with altered implementations of numerosity representations (x_sample_ + x_distractor_ + x_SD interaction_) for the model to fit. The temporal part of the demixed data was unchanged. Three different implementations were created: first, a non-structured Gaussian distribution of neuronal loadings and no alignment to any components (cp. **Fig. 1d**); second, a distribution with the same degree of sparsity as the original data, but with sparse axes randomly rotated to align to other components (cp. **Fig. 1e**); third, a substitute with the same sparse distribution of neuronal loadings as in the original data (cp. **Fig. 1f**).

The model captured an increasing proportion of variance of the full signal across the three substitutes (p < 0.001; one-way ANOVA; **Fig. 7c**). The absolute differences in explained variance were comparatively small (left axis), but remarkable in relation to the variance of the manipulated signal (right axis) and given that the representational geometry was unchanged and identical for all substitutes (cp. **Fig. 1**). A comparable result was obtained for the explained variance of the numerosity coding part (p < 0.001; one-way ANOVA; **Fig. 7d**).

Taken together, these results demonstrate that sparse implementations of working memory representations are favored by recurrent circuits, the characteristic wiring motif of association cortices such as the PFC.

## Discussion

We presented a framework to examine the contributions of individual neurons to population-level responses in representation space and to utilize its implementation structure. We identified heavy-tailed, i.e., sparse distributions of neuronal loadings on components that captured disentangled and sequential memory representations including the recovery of memory content after distraction. The switching of working memory components circumvented interference. These components could be traced to small subpopulations of neurons with distinct electrophysiological properties and temporal dynamics. Modelling showed that such sparse implementations with sequentially active components are supported by recurrently connected networks.

### Bridging population activity and neuronal implementation

Population-level activity and representational geometry were previously studied without forming direct links to individual neurons (Bernardi et al., 2020; Chung & Abbott, 2021; Kriegeskorte & Wei, 2021; Okazawa et al., 2021). However, while single-neuron selectivity measures have the advantage of being more easily connected to biological properties such as cell type, receptor expression and axonal projection targets, they are typically chosen based on intuition and past experience and only partially or indirectly reflect the full representational space (Hirokawa et al., 2019; Jacob & Nieder, 2014).

Our sparse component analysis (SCA) framework (**Fig.1**) combines the advantages of both perspectives. It builds on representational geometry for a comprehensive account of the data and then links the relevant coding dimensions in the activity space to populations of strongly contributing neurons, which allows relating the population-wide activity patterns to tangible physiological measures.

### Implementation reveals biologically relevant dimensions in activity space

Without respecting implementation, selecting components in activity space for further analysis is arbitrary. It is often done post-hoc after visualizing the top PCs, or by relying on the heuristics of ‘what should be coded’ in the system (Aoi et al., 2020; Bernardi et al., 2020; Libby & Buschman, 2021). This approach becomes problematic when the dimensionality is too high or when too many variables are involved.

By exploiting neuronal implementation, SCA identifies activity components in an un-biased and non-arbitrary way. SCA can therefore capture a more complete set of stimulus-associated variables (dimensions), most notably the temporal modulation of stimulus coding. This reduces bias otherwise introduced by selecting specific time windows, across which neuronal activity is averaged, and acknowledges the role of different response dynamics for information coding (Bondanelli & Ostojic, 2020; Mante et al., 2013). Furthermore, incorporating temporal modulation renders analyses more robust to noise (Johnstone & Lu, 2009), which is usually Gaussian and could hide the structure in implementation.

The implementation’s sparse structure is a result of biological constraints regarding the connections among individual neurons. The approximately 10^4^ dendritic spines on each cortical neuron (Eyal et al., 2018) define an upper limit for the number of neurons it could read out from. The 10^9^ neurons in a cortical region such as human PFC (Courchesne et al., 2011; Herculano-Houzel et al., 2015), and even sub-modules with one to two magnitudes fewer neurons, therefore cannot be reached directly. The addition of one connection step would allow reaching the majority of PFC neurons, but at the cost of producing a layer of 10^4^ to 10^5^neurons that are dedicated exclusively to feeding the single hypothetical downstream neuron. This is prohibitively inefficient. In such polysynaptic chains, it is more likely that meaningful representations have already emerged in intermediate layers as a result of direct connections from the source region. This notion is also in line with the high dimensionality and non-linear mixed selectivity characteristic of PFC, which allow for direct linear readout of complex representations without further computations (Rigotti et al., 2013).

Neurons share inputs and have local recurrent connections, which are particularly pronounced in association cortices such as the PFC (Harris et al., 2019), resulting in more similar firing patterns among neurons within cortical regions. Consequently, neurons might display activity that is weakly correlated to some components of the representational geometry even though they do not participate in the readout. This emphasizes the importance of truncating neurons with weak loadings and enforcing sparsity constraints for estimating potential readout connections (**Fig. 4**) and motivates the use of dynamical systems modelling to validate correlative measures (**Fig. 7**).

### Working memory persistence without neuronal persistence

Applied to working memory maintenance in the face of distraction, our framework uncovered an unexpected sequential representation of numerosity information across multiple task epochs (**Fig. 2**). This result was neither encouraged nor guaranteed by SCA. This suggests that the readout of memory content from the PFC is optimized for accuracy in each behavioral context rather than optimized for stability across time periods. The distractor occupied the same resources as the sample numerosity with regard to the sensory and memory component (**Fig. 3**), forcing behaviorally relevant information to be shifted to the recovery component following distraction. Thus, working memory content was maintained by distinct mechanisms before and after interference (**Fig. 4**).

The subpopulation of recovery neurons was characterized by electrophysiological properties that set these neurons apart from the other populations and could render them particularly suited to working memory storage. Their longer intrinsic timescales (**Fig. 5**) suggest more stable memory retention (Kim & Sejnowski, 2021; Murray et al., 2014). These neurons also distinguished between sample and distractor contexts, which is crucial for determining what information to keep and what information to discard (**Fig. 6**). The contextual signal was additively mixed with the numerosity coding signal in these neurons, but might still act as gain modulation for numerosity information given the neuronal input-output non-linearity (Dubreuil et al., 2020).

Representing memory content by sequentially active subpopulations is advantageous. With relay of information, a result of locally feed-forward connectivity, a network can maintain multiple inputs from previous time points and show more resistance to noise (Orhan & Pitkow, 2020). Furthermore, the PFC might be non-linearly mixing context and memory representations in all possible ways, expanding dimensionality to enable flexible readout (Rigotti et al., 2013). Extensive training could have strengthened the non-linear mixture of second memory epoch context and sample numerosity representations that was most important in the current task, with the PFC retaining other mixtures (e.g. the component coding for sample numerosity in the first memory epoch) for other behavioral demands. In this view, the subpopulation of memory neurons could function as a more passive short-term memory storage oblivious to the behavioral relevance of the memorized information.

Introducing distraction into the memory delay unmasked the crucial role of recovery neurons for working memory maintenance, which would have been hidden in simpler tasks. This highlights the importance of including richer temporal structure, multiple processing stages and behavioral perturbation into cognitive task designs to enable dissection of higher-order brain functions in finer detail and sampling from the full spectrum of underlying mechanisms.

### Alternative implementation structures

We focused here on detecting sparse structure in the representational geometry’s neuronal implementation, which is linked to the standardized moment of kurtosis. Consequently, the loading distributions have both positive and negative heavy tails. Reading out a given sparse component thus requires both excitatory and inhibitory connections. However, long-range corticocortical projections are mainly excitatory. This means that other selection criteria that capture non-symmetrical structure such as the standardized moment of skewness should also be explored (Koren et al., 2020; Román Rosón et al., 2019).

Structure could be in the form of disjointed cell clusters (Hirokawa et al., 2019) or a mixture of Gaussians (Dubreuil et al., 2020). However, if present, these structures would not dissect the representational geometry, as they do not have a one-to-one relation to the dimensions in the activity space. Our neuronal implementation followed a unimodal Laplace distribution (Fig. 2g) instead of a multimodal distribution.

Structure can also be investigated when there are no prior assumptions about the underlying distributions of neuronal loadings. For example, given that neuronal firing is energy-consuming and non-negative, possibly encouraging neurons to align to the dimensions of the representational geometry that have shorter ranges of variation, non-uniform distributions of the number of selective neurons across different dimensions can arise (Whittington et al., 2022). However, because all neurons are counted equally, structure probed non-parametrically could potentially be clouded by the large number of weakly coding (non-dominant) neurons and thus difficult to detect, in particular in PFC (Bernardi et al., 2020).

### Relation of SCA to other linear dimensionality reduction methods

Different linear dimensionality reduction methods based on L2 reconstruction loss will yield comparable representational geometries, but they will not find the same projections of the representational geometry, i.e., the same components or the same coordinate system in which the data is expressed. The principle components of PCA are conveniently orthogonal and ranked by variance (Vu & Lei, 2013), but usually neither correspond to task-related components nor align to the activity of individual neurons (Higgins et al., 2021). Truncating the smaller PCs provides denoised signal as a preprocessing step for independent component analysis (ICA) that can infer the independent sources in the signal space (Hyvärinen & Oja, 2000). Its most common form, fastICA, enforces sparsity constraints on the activity of the components, reflecting an assumption about the activity (Hyvarinen, 1999). In contrast, in SCA the sparsity constraint is on the neuronal implementation, i.e., the potential readout weights corresponding to the mixing matrix in ICA, reflecting an assumption about the connectivity.

Neuronal representations must be communicated. Information that cannot be accessed by other neurons does not exist. In order to understand complex neural systems such as the PFC where we lack clear priors about the signal sources, it is paramount to exploit the circuit and wiring motifs that underlie the observed activity patterns.

## Acknowledgements

This study was supported by grants from the German Research Foundation (DFG JA 1999/1-1, JA 1999/5-1, JA 1999/6-1) and the European Research Council (ERC StG MEMCIRCUIT, GA 758032) to S.N.J and grants NI 618/10-1 and NI 618/13-1 to A.N.

## Author contributions

X.-X.L. conceived the study and performed the analyses with contributions from S.N.J. A.N. and S.N.J. designed the experiments and collected the data. X.-X.L. and S.N.J. wrote the manuscript and prepared the figures. All authors edited the manuscript.

## Declaration of interests

The authors declare no competing interests.

## Methods

Two adult male rhesus monkeys (*Macaca mulatta*, 12 and 13 years old) were used for this study. All experimental procedures were in accordance with the guidelines for animal experimentation approved by the national authority, the Regierungspräsidium Tübingen. A detailed description is provided elsewhere (Jacob et al., 2018; Jacob & Nieder, 2014).

### Surgical procedures

Monkeys were implanted with two right-hemispheric recording chambers centered over the principal sulcus of the lateral prefrontal cortex (PFC) and the ventral intraparietal area (VIP) in the fundus of the intraparietal sulcus. This study reports on the PFC data.

### Task and stimuli

The animals grabbed a bar to initiate a trial and maintained eye fixation (ISCAN, Woburn, MA) within 1.75°of visual angle of a central white dot. Stimuli were presented on a centrally placed gray circular background subtending 5.4° of visual angle. Following a 500 ms pre-sample (pure fixation) period, a 500 ms sample stimulus containing 1 to 4 dots was shown. The monkeys had to memorize the sample numerosity for 2,500 ms and compare it to the number of dots (1 to 4) presented in a 1,000 ms test stimulus. Test stimuli were marked by a red ring surrounding the background circle. If the numerosities matched (50 % of trials), the animals released the bar (correct Match trial). If the numerosities were different (50 % of trials), the animals continued to hold the bar until the matching number was presented in the subsequent image (correct Non-match trial). Match and non-match trials were pseudo-randomly intermixed. Correct trials were rewarded with a drop of water. In 80 % of trials, a 500 ms interfering numerosity of equal numerical range was presented between the sample and test stimulus. The interfering numerosity was independent from either the sample or test numerosity and therefore not useful for solving the task. In 20 % of trials, a 500 ms gray background circle without dots was presented instead of an interfering stimulus, i.e., trial length remained constant (control condition, blank). Trials with and without interfering numerosities were pseudo-randomly intermixed. Stimulus presentation was balanced: a given sample was followed by all interfering numerosities with equal frequency, and vice versa. Throughout the monkeys’ training on the distractor task, there was never a condition where a stimulus appearing at the time of the distractor was task-relevant.

Low-level, non-numerical visual features could not systematically influence task performance (Jacob & Nieder, 2014; Nieder et al., 2002):in half of the trials, dot diameters were selected at random. In the other half, dot density and total occupied area were equated across stimuli. CORTEX software (NIMH, Bethesda, MD) was used for experimental control and behavioral data acquisition. New stimuli were generated before each recording session to ensure that the animals did not memorize stimulus sequences.

### Electrophysiology

Up to eight 1 MΩ glass-insulated tungsten electrodes (Alpha Omega, Israel) per chamber and session were acutely inserted through an intact dura with 1 mm spacing. Single units were recorded at random; no attempt was made to preselect for particular response properties (Jacob & Nieder, 2014). Signal amplification, filtering, and digitalization were accomplished with the MAP system (Plexon, Dallas, TX). Waveform separation was performed offline (Plexon Offline Sorter).

### Data analysis

Data analysis was performed with Python using custom scripts based on packages NumPy, SciPy, sci-kit learn, TensorFlow2, PyTorch, Matplotlib and Plotly.

### Preprocessing

Single units were included in the analysis if they were recorded in at least 4 correct trials of each task condition (meaning each unique sample and distractor numerosity combination). This resulted in 467 neurons across 78 sessions recorded in the PFC. Trials without distractors were not included in the analyses unless specified otherwise.

Unless specified otherwise, the firing rates were binned in a Gaussian window with sigma of 50 ms and step of 100 ms, aligned to the start of the fixation period. The data were then organized into a neuron-by-condition-by-timepoint tensor. Each tensor entry was normalized by the standard deviation across trials (within each condition).

### Demixing

Given the independence of the task variables sample numerosity (s), distractor numerosity (d) and trial time (t), the neuronal activity can be directly factorized into parts for each variable and their interaction:

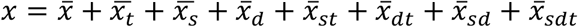

Because the stimulus response is also modulated by time, each part was grouped together with its interaction with time (Kobak et al., 2016):

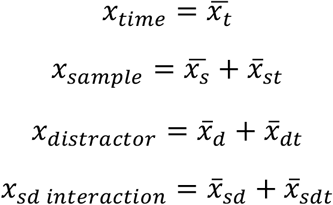

### Visualization of representation and implementation space

For a data matrix *X* where each column vector *x* is the demixed activity of a neuron, the singular value decomposition was taken:

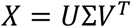

where *U* and *V* are unitary matrices and Σ is a diagonal matrix with ordered singular values. The first *n* columns of *U*Σ are the PCs that were used to visualize the representational geometry. The first n columns of *V*Σ are loadings on the PCs that were used to visualize the implementation space.

Within this subspace an arbitrary component can be specified with *UΣP*_:,1_ (*P*_:,1_ being a column vector from a unitary matrix *P*), with the orientation of this component given by *P*_:,1_. The loadings on this component will be the first row of (*UΣP*)^+^ *X*= *P*^*T*^*V*^*T*^, that is *P*_:,1_^*T*^*V*^*T*^. This way, the loadings are visualized with the same orientation *P*_:,1_. in implementation space as their corresponding component in representation space. The sparsity index of the neuronal loadings on component *UΣP*_:,1_ is then:

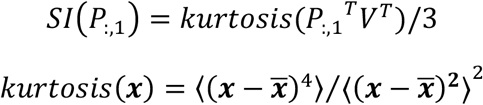

### Sparse component analysis

Following the formulation of sparse coding (Georgiev et al., 2007; Lee et al., 2007; Olshausen & Field, 1996), sparse component analysis (SCA) reduces the dimensionality of the dataset and extracts the unique components by enforcing a sparse penalty on neuronal loadings:

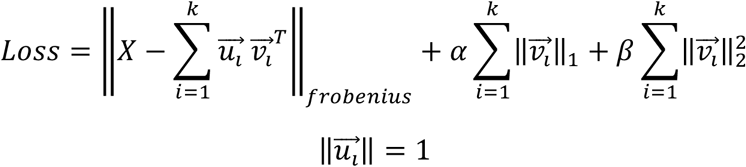

The loss function is defined as the sum of the reconstruction loss and the regularizations. Data *X* is organized as a *n* firing instances by *p* neurons matrix. *X* is then approximated by *k* firing activity vectors 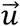 and their corresponding neuronal loadings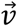. Parameter α controls the strength of L1-regularization that encourages sparsity of the loadings. Parameters α and *k* were determined by a cross-validated grid search. β was set at 0.01 to smooth the loss landscape and make the result stable across random initializations.

### Substitute data for SCA

Substitute data were created for the demixed sample coding part *X* of the data (Fig. 2). For the singular value decomposition *X*= *UΣV*^*T*^, *U*Σ specifies the representational geometry (see above). Operations were performed on *V* only.

A random unitary matrix *R* with the size of the number of neurons was drawn from a Haar distribution. The original matrix *V* was replaced with *V*′ = *VR. V*′ is also a unitary matrix, meaning that this manipulation will not change the geometries but will rotate them to random axes. In other words, it will linearly combine the loadings including those on the components with very low variance, which will render the substitute distribution of loadings on the sample numerosity components close to Gaussian. The substitute data is then *X*′ = *UΣV*′^*T*^ = *XR*

### Measures of sparse component activity

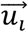 in SCA specifies the activity of the sparse component *i*. The following measures of the set of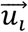were compared between the original dataset and its substitutes (n = 1000).

#### Spread of representation

The standard deviation of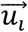 across different numerosity conditions *k* at each time point was used to define the relative (normalized) information at that time point. Specifically, each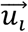 was first reshaped into a condition-by-timepoint matrix *Y*^*i*^. Then the information in component *i* at time point *t* is given by:

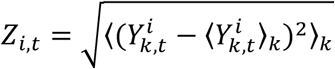

The skewness of the information across time points was calculated for each component and averaged across components as follows:

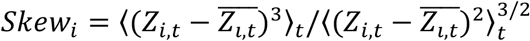

Positively skewed *Z* indicates a long tail in the distribution of information across time points, corresponding to few time points having high information. Conversely, a smaller or even negative skewness implies there are more high information timepoints than low information time points, making the high information more spread out across time points. We define the spread of representation as the negative skewness:

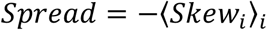

#### Overlap of active periods

The dot product of the information of every pair of components *i* and *j* was taken and averaged across pairs:

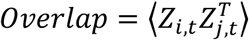

#### Maximum tuning reversal

A given component *i* may show changes of tuning to sample numerosities during the course of a trial. Its tuning at time *t* is specified by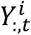. For each component *i*, the dot product similarity of tunings between timepoint pairs was specified in the non-diagonal entries in 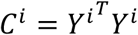 where the diagonal entries are the strength of the tuning at each time point. *C*^*i*^ was then normalized to the strongest tuning: 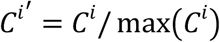 The most negative entry in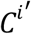 was then the degree of reversal in this component. 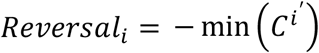. It would reach the maximum of 1 when tuning at a given time point is the complete reversal of the strongest tuning. It would be close to 0 when the tuning does not reverse. The maximum tuning reversal is then the largest reversal in a set of SCs:

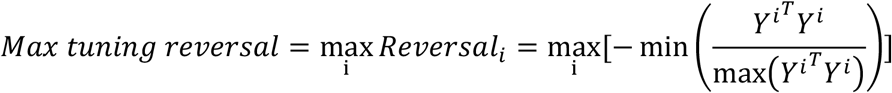

#### Component similarity

Let *U*_*sca*_ be the concatenation of activity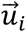 and *V*_*sca*_ the concatenation of loadings 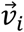 of the sparse component *i*. The data matrix can be expressed as 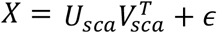. *ϵ* denotes the noise term. Then it follows 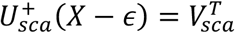. The pseudoinverse 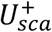 can be viewed as a linear transform of the original data. Since all the activities 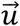 have unit length, larger loadings would be required to express an arbitrary geometry when the activities are correlated, meaning lower efficiency. The component similarity is measured by the product of the singular values of *U*_*sca*_. Formally, if the singular value decomposition gives *U*_*sca*_= *UΣV*^*T*^, then

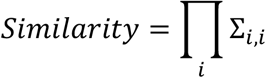

The similarity can also be viewed as the determinant of the transformation matrix from arbitrary orthogonal bases to the bases of *U* _*sca*_

### Numerosity information in different components

The standard deviation *Z*_*i,t*_ for all time points *t* specifies the evolution of normalized information within this component. But since 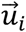 in component *i* has unit length, this measure does not allow for direct comparisons between components (see above). To allow for such comparisons (Fig. S1), the norm of 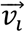 is therefore applied to *Z*_*i,t*_ as a scaling factor:

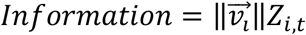

### Linear discriminant analysis decoding

Neurons recorded in different sessions were stitched together. To account for the different number of trials recorded per neuron, a criterion was set to ensure there were at least 1.5 times more trials than neurons. This resulted in 228 neurons with at least 385 trials each. Removing incorrect trials and selecting the minimum number of trials recorded per condition and neuron left 118 trials per neuron. Trials of the same condition were then randomly selected for each repetition of the analysis.

Multi-class linear discriminant analysis (LDA; sci-kit learn package) was used for decoding because of its advantageous property of accounting for data covariance. LDA assumes the same covariance in every class. It finds the projection that preserves the Mahalanobis distance between classes and predicts the label of a new data point by its Mahalanobis distance to the class centroid. Shrinkage of the measured covariance matrix was performed by averaging with a diagonal matrix. The strength of shrinkage was determined following the Ledoit-Wolf lemma (Ledoit & Wolf, 2004).

Decoding accuracy, i.e., the ratio of correctly predicted trials, was averaged across 7 repetitions of 7-fold cross-validation.

### Spike train statistics

Firing rates were binned in a Gaussian window with sigma of 12.5 ms and step of 25 ms.

Correlation, autocorrelation and intrinsic timescales were determined as described elsewhere (Murray et al., 2014). The firing rate of each neuron *n* at timepoint *t* of trial *i* is expressed as *x*_*n,i,t*_. The Pearson correlation between timepoints *t*1 and *t*2 is then:

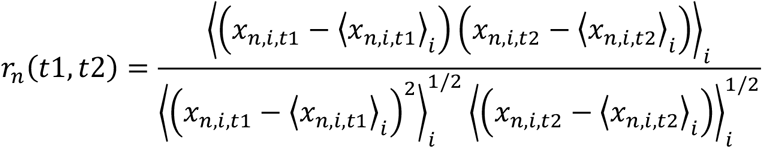

Autocorrelation is defined as:

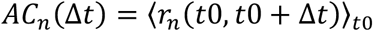

To account for the refractoriness and adaptation at small time lags, fitting started at the time lag where the autocorrelation function had dropped most strongly. Neurons with the strongest drop after 400 ms were discarded (6 neurons). The autocorrelation was then fitted with an exponential decay:

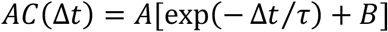

Parameters *A* and *B* were constrained in [0,1] and *τ* was constrained from 10 ms to 2000 ms. The autocorrelation function of 8 neurons could not be fitted. The neurons with *τ* fitted below 20 ms (20 neurons) or above 1600 ms (25 neurons) were excluded because of the biologically unrealistic fit. This left 408 neurons. Very few neurons were excluded in the dominant subpopulations (2, 2, and 1 neurons for the sensory, memory and recovery subpopulation, respectively).

The inter-spike intervals (ISI) were determined for the entire session. The coefficient of variation (CV) measures the global variation of a neuron’s ISI and is defined as:

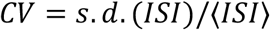

In contrast to CV, local variation (LV) measures the local ISI change (Shinomoto et al., 2009). It is defined as:

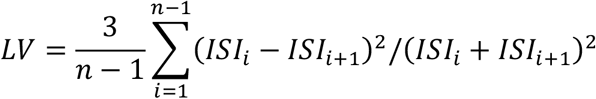

CV and LV are both expected to be 1 for spiking activity following a Poisson process. CV and LV would be 0 for perfectly regular firing and larger than 1 for more irregular firing than by a Poisson process.

### Kullback-Leibler divergence

KL divergence measures the difference between two distributions. For the analyses of intrinsic time scales and periodicity, KL divergence was calculated between the distribution of statistic *x* for the entire population *P* and that of sub-samples *Q* (either dominant subpopulations or bootstrap subsamples). It is given by:

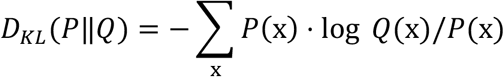

To create the null distribution of *D*_*KL*_, 27 neurons (comparable to the number of neurons in the dominant subpopulations after exclusion of neurons in which no autocorrelation function could be fitted) were randomly sampled from the PFC population 1000 times.

### Temporal dynamics

#### Periodicity

The Fourier transform of the demixed temporal part of the firing rate of each neuron is given by:

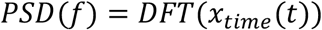

Then, the periodicity was defined as the ratio between the power of the harmonics of 1/1.5 Hz (reflecting the onset of visual input at regular spacing of 1.5 s) and the power of all frequencies:

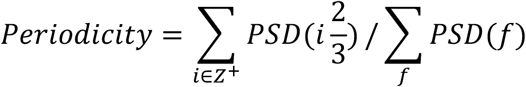

#### Tangling

Tangling reflects the smoothness and stability of the flow field around the vicinity of state *x*_*t*_ on a trajectory (Russo et al., 2018). It is given by:

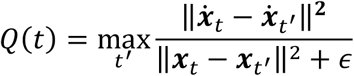

It specifies the maximum difference between the derivative at state *x*_*t*_ and the derivative at other states ***x***_*t*′_, normalized by their Euclidean distance. A small constant *ϵ* was added to avoid numerical error when the two states were too close.

### Recurrent neural network

A recurrent neural network (RNN) model was implemented using the PyTorch neural network module. The model has the formulation:

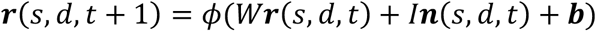

***r*** is the firing rate of units in the condition of sample numerosity *s* and distractor numerosity *d* at time point *t. ϕ* is the non-linear activation function, chosen to be a rectified linear unit (ReLu) to respect the biological characteristics of non-negative firing rates with high upper limits. *W* is the within-population connectivity matrix. *I* is the input matrix with the dimensions of 467 (total number of units) by 4 (number of numerosities). A column *I*_:,*a*_ is the input to the units when numerosity *a* is being presented. ***n*** is an indicator vector with the entry ***n***_*a*_ corresponding to the presented numerosity being 1 and all other entries being 0. ***b*** is the intercept. *W, I* and ***b*** are the parameters to be trained. Formally, ***n*** as a function of trial type specified by *s* and *d* and time point *t* is defined by:

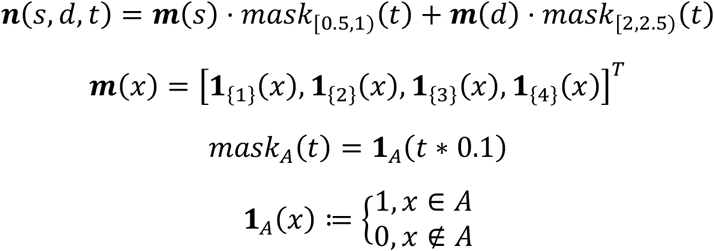

***m*** maps a numerosity to the corresponding one-hot vector. *mask*_*A*_(*t*) indicates the time (0.1 s steps) when the corresponding stimulus is presented. **1**_*A*_(*x*) is an ancillary indicator function to define ***m*** and *mask*.

The model was trained to produce the whole sequence of firing rates ***r***(*s, d, t*) in order to match the target data ***x***_*s,d,t*,_ given the initial firing rate in the fixation period ***r***(*s, d*, 0) and the input ***n***(*s, d, t*). The loss function is defined as:

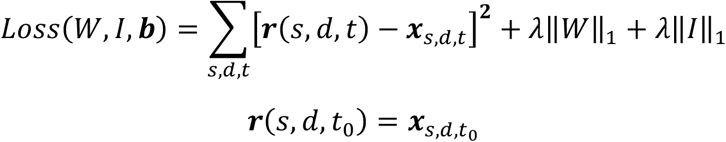

The coefficient *λ* controls the strength of regularization and was determined by a grid search with cross validation.

The prediction of the later timepoints relies on the quality of the prediction of the early timepoints. If the training was done only by giving the first timepoint, convergence would be difficult to achieve and learning heavily biased towards reproducing early timepoints in the data. To overcome this possible instability, the model was trained in a recursive fashion by first using every timepoint as the initial firing rate, training the model to predict the following timepoints and gradually increasing the number of timepoints the model needs to predict. As such, at each iteration *i*, the temporal sequence *x*_*s,d,t*_ was reorganized into *T* − *i* chunks of length 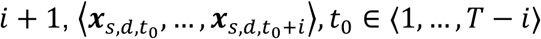, with the first firing rate in each chunk as initial firing rate and the rest as target to be fit by the model.

### Variance explained by RNN

The variance explained by the model was determined by the difference between the model’s predicted trajectory and the trajectory of the original data normalized to the difference between a reference trajectory (constant activity set to the first entry of the fixation period) and the trajectory of the original data:

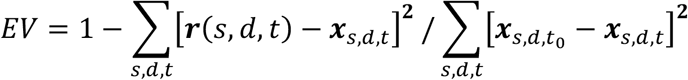

The normalized EV (Fig. 7c, right axis) was defined as the difference between a substitute’s EV and the original data’s EV, divided by the percentage of the manipulated variance (numerosity coding signal, 27.4 %; cp. Fig. 2b). EV for the numerosity signal (Fig. 7d) was calculated by replacing both ***r***(*s, d, t*) and ***x***_*s,d,t*_ with their demixed numerosity representing parts.

### Substitute data for RNN

In order not to distort the strong connection between sample and distractor numerosity coding (e.g., Fig. 3b, Fig. S1), the loadings of these two parts of the data and their interaction were shuffled together to create three types of substitute datasets. The RNN model was then trained on the substitutes.

#### Gaussian distribution of loadings

The Gaussian substitutes were created as described for SCA, except for that singular value decomposition was performed on *X*_*sample*_ + *X*_*distractor*_ + *X*_*sd_interaction*_ = *X*_*all*_− *X*_*t*_ = *UΣV*^*T*^.

#### Sparse distribution with random alignment

For *k* dimensions of the numerosity coding part of the data (determined by cross validation), a *k* × *k* unitary matrix *R* was randomly drawn from a Haar distribution and combined with an identity matrix *I* to create 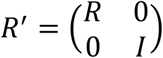. Then, *V*′ = *VR*′ was substituted for *V*. This leaves the sparse structure in the original *k* dimensional numerosity representing subspace intact, but rotates the sparse structure in *V*_:,1:*k*_ to random orientations.

#### Sparse distribution with original alignment

The rows of *V*_:,1*:k*_, i.e., the neuronal identities, were permuted by substituting *V*′ = (*V*_*permute*,1*:k*_, *V*_*:k*+1*:p*_) for *V*.

**Fig. S1.**
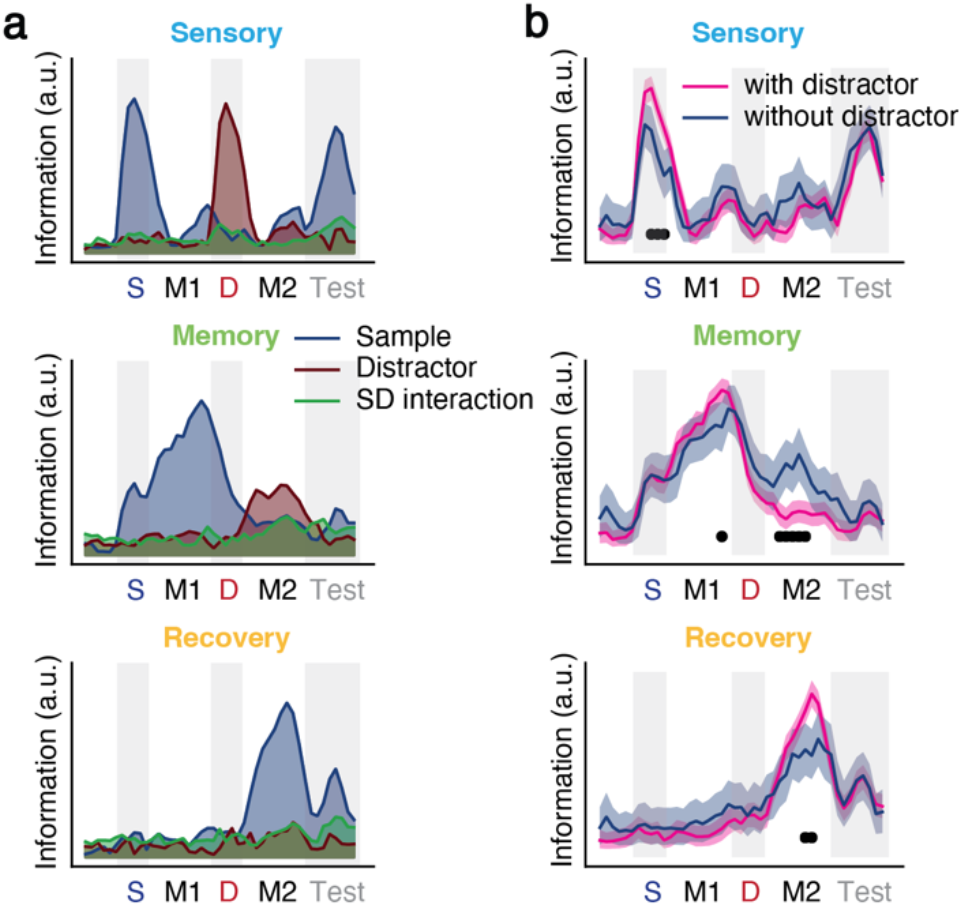
The effect of distraction on sample numerosity sparse components. (**a**) Information (standard deviation across conditions) about sample numerosity, distractor numerosity and their interaction in each of the three sample numerosity sparse components (SCs) in trials with a distractor. (**b**) Sample numerosity information as in (a) for the three SCs in trials with and without a distractor. Shaded area indicates [2.5 %, 97.5 %] confidence interval. Black dots indicate timepoints with significant differences (p < 0.00125, bootstrap).

**Fig. S2.**
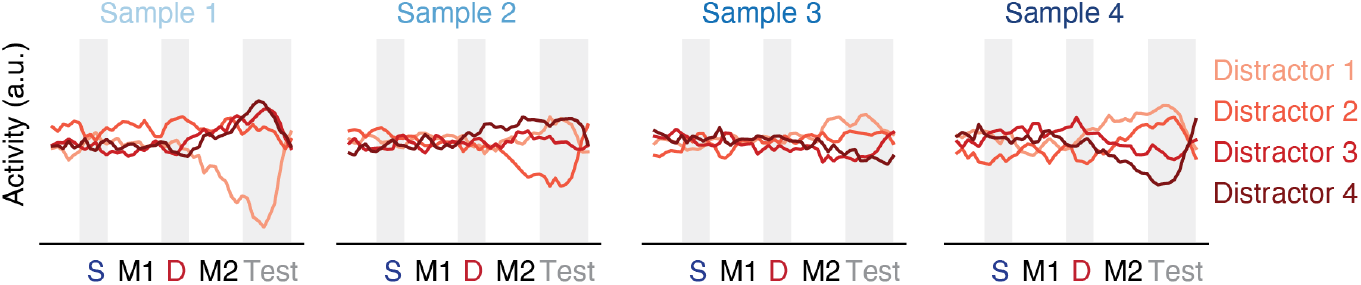
Sample-distractor interaction sparse component. SCA performed on the demixed sample-distractor interaction part of the data identified one component that optimally reconstructed the data using cross-validation. The activity of this SC is shown for all sample-distractor combinations.

## Notes

### Competing Interest Statement

The authors have declared no competing interest.

